# Geographic distribution of feather *δ*^34^S in Europe

**DOI:** 10.1101/2023.03.10.532116

**Authors:** Vojtěch Brlík, Petr Procházka, Luana Bontempo, Federica Camin, Frédéric Jiguet, Gergely Osváth, Michael B. Wunder, Rebecca L. Powell

**Affiliations:** Department of Ecology, Charles University, Prague, Czech Republic; Czech Academy of Sciences, Institute of Vertebrate Biology, Brno, Czech Republic; Traceability Unit, Research and Innovation Centre, Fondazione Edmund Mach, Trento, Italy; Center Agriculture Food Environment (C3A), University of Trento, Via E. Mach, 1 - 38010 38098 San Michele all’Adige (TN), Italy; International Atomic Energy Agency, Vienna International Centre, PO Box 100, A-1400 Vienna, Austria; Museum National d’Histoire Naturelle, Sorbonne Université, CNRS, UMR7204 CESCO, Paris, France; Musem of Zoology, Academic Cultural Heritage Department, Babeş-Bolyai University, Cluj-Napoca, Romania; Evolutionary Ecology Group, Hungarian Department of Biology and Ecology, Babeş-Bolyai University, Cluj-Napoca, Romania; University of Colorado Denver, Department of Integrative Biology, Denver, USA; University of Denver, Department of Geography and the Environment, Denver, USA

**Keywords:** feather, isoscape, sulphur, wetland

## Abstract

Geographic distribution models of environmental stable isotopes (so called ‘isoscapes’) are widely employed in animal ecology, wildlife forensics and conservation. However, the application of isoscapes is limited to elements and regions for which the spatial patterns have been estimated. Here, we focused on the ubiquitous yet less commonly used stable sulfur isotopes (*δ*^34^S). To predict the European *δ*^34^S isoscape, we used 242 feather samples from Eurasian Reed Warbler (*Acrocephalus scirpaceus*) formed at 69 European wetland sites. We quantified the relationships between sample *δ*^34^S and environmental covariates using a random forest regression model and applied the model to predict the geographic distribution of *δ*^34^S. We also quantified within-site variation in *δ*^34^S and complementarity with other isotopes on both individual and isoscape levels. The predicted feather *δ*^34^S isoscape shows only slight differences between the central and southern parts of Europe while the coastal regions were most enriched in ^34^S. The most important covariates of *δ*^34^S were distance to coastline, surface elevation and atmospheric concentrations of SO_2_ gases. The absence of a systematic spatial pattern impedes the application of the *δ*^34^S isoscape but high complementarity with other isoscapes advocates the combination of multiple isoscapes to increase precision of animal tracing. Feather *δ*^34^S compositions showed considerable within-site variation with highest values in inland parts of Europe, likely attributed to wetland anaerobic conditions and redox sensitivity of sulfur. The complex European geography and topography as well as using *δ*^34^S samples from wetlands may contribute to the absence of a systematic spatial gradient of *δ*^34^S values in Europe. We thus encourage future studies to focus on the geographic distribution of *δ*^34^S using tissues from diverse taxa collected in various habitats over large land masses in the world (i.e., Africa, South America or East Asia).

**Open Research statement:** Sample metadata are stored and available for reuse in the AviSample Network database (https://avisample.net/: AS00001–AS00117; Brlík et al. 2022b). The information on the isotopic composition and geographic origin of feather samples used to produce the *δ*^34^S isoscape, predicted feather *δ*^34^S, *δ*^2^H and *δ*^13^C isoscapes are publicly available from the Zenodo data repository at https://doi.org/10.5281/zenodo.7315567 (Brlík et al. 2022c).

## Introduction

The stable isotopic compositions of ecosystem components vary spatially due to a wide variety of biogeochemical processes (West et al. 2010). These geographic distributions of isotopes (i.e., ‘isoscapes’) can be used to infer spatial origin and have manifold applications, especially in animal ecology, wildlife forensics and conservation (Hobson and Wassenaar 2018). Isoscapes are used to infer geographic space use and thus applied to study altitudinal migration (Villegas et al. 2016), to quantify strength of migratory connectivity (Norris et al. 2006) or to trace natal origins of animals (Procházka et al. 2013; Brlík et al. 2022a). Geographic space use can affect animals and stable isotopes thus may indirectly inform demographic models to understand population changes and target conservation efforts (Jiguet et al. 2019). Continental-scale isoscapes essential for tracing origins are currently available for *δ*^2^H, *δ*^13^C, *δ*^15^N, *δ*^18^O and ^87^Sr/^86^Sr, but their application is limited to regions with steep isotopic gradients or isotopically-distinct regions (Amundson et al. 2003, West et al. 2010, Bataille et al. 2020).

Some continental and global isoscapes feature systematic spatial patterns on continental scales (e.g., *δ*^2^H and *δ*^18^O in North America and East Asia; Bowen and Revenaugh 2003) while others show weak spatial structure (e.g., *δ*^13^C and *δ*^15^N in Africa and South America; Amundson et al. 2003, Powell and Still 2010) limiting their application. Alternatively, the combination of spatial predictions using multiple isoscapes can be used to increase precision of tracing (e.g., Popa-Lisseanu et al. 2012, Veen et al. 2014). Either way, developing isotopic maps is essential for broadening the potential of isotopes in animal ecology and wildlife forensics.

Geographic distributions of stable sulfur isotopes (*δ*^34^S) in recent organic tissues have already been modelled for Great Britain, Tanzania, sub-Saharan Africa, and the conterminous USA (Valenzuela et al. 2011, Kabalika et al. 2020, Newton 2021, Brlík et al. 2022a). Additionally, a recent study used archaeological remains to predict the geographic distribution of *δ*^34^S in Europe (Bataille et al. 2021). Moreover, deposition of marine-derived sulphates from coastal towards inland areas was identified as the most important driver of *δ*^34^S geographic distribution (Lott et al. 2003, Zazzo et al. 2011). Gradients formed by the deposition of marine sulphates have been detected on both regional (Wadleigh and Blake 1999, Zazzo et al. 2011) and continental scales (Valenzuela et al. 2011, Bataille et al. 2021; Brlík et al. 2022a). In addition, spatial variation in atmospheric concentrations of combustion gases and bedrock geochemistry, have also been found to affect spatial patterns of *δ*^34^S values (Case and Krouse 1980, Thode 1991). Indeed, detailed knowledge of these relationships could be used to predict *δ*^34^S geographic distributions on continental scales. The isotopic offset between environment and organic tissues is generally low for *δ*^34^S (Felicetti et al. 2003, McCutchan et al. 2003, Florin et al. 2011, Tcherkez and Tea 2013) providing an opportunity to apply a single *δ*^34^S isoscape to a range of taxa. Despite these benefits, continental *δ*^34^S isoscapes of organic tissues are currently available only for part of North America and for human archaeological remains in Europe (Valenzuela et al. 2011, Bataille et al. 2021).

In this study, we (i) develop a European *δ*^34^S isoscape using feather keratin samples, (ii) quantify within-site variation in *δ*^34^S, and (iii) quantify complementarity of *δ*^34^S with other feather isotopes (individual level) and other isoscapes (spatial level).

## Methods

### Sample collection

We used 242 feathers from Eurasian Reed Warbler (*Acrocephalus scirpaceus*) with known geographic origin at 69 European wetland sites previously used in Procházka et al. (2013) and newly collected in 2019 (Fig 1). At these sites, we collected the innermost primary feathers from one to four juvenile Eurasian Reed Warblers (one sample = 8 sites; two = 2; three = 6 and four = 53) during the fledgling period (June-early August; Fig 1). Juveniles of the Eurasian Reed Warbler are mostly stationary during this period before post-fledging dispersal (Mukhin 2004, Grinkevich et al. 2009) and we consider the sampling locations their natal origins and thus known geographic origins of these samples.

**Fig 1.**
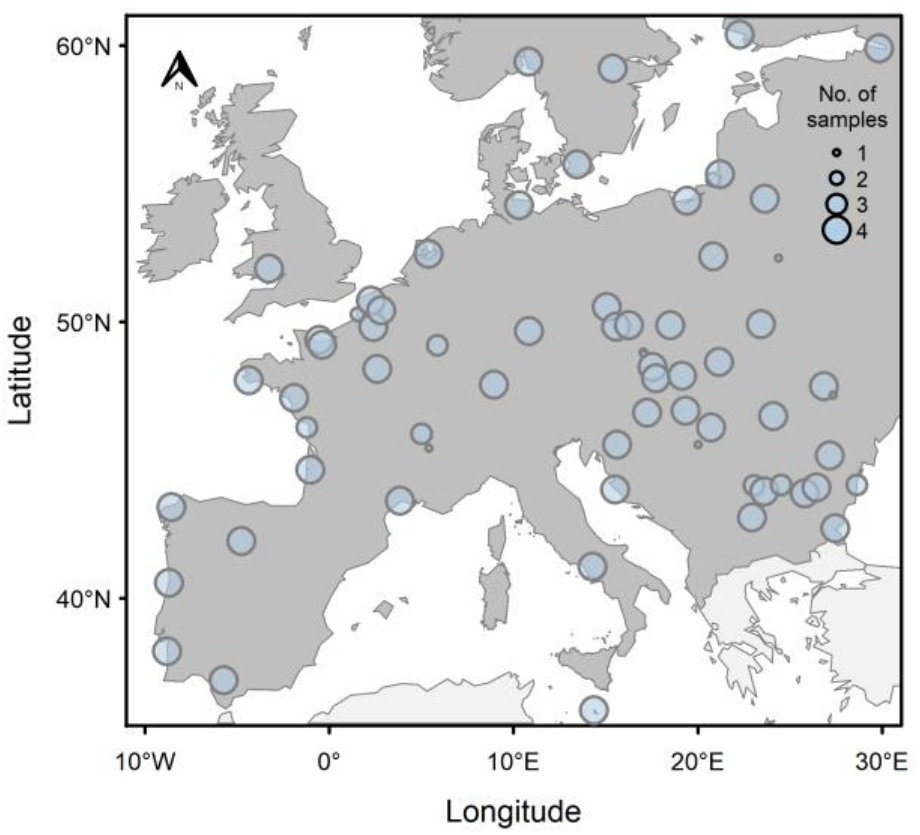
Locations of European sampling sites (n = 69) coinciding with the geographic origins of feather samples collected from juvenile Eurasian Reed Warblers (Acrocephalus scirpaceus; n = 242). Dark grey region depicts the extent of prediction.

### Isotopic analysis

We cleaned the feather samples in a 2:1 chloroform:methanol mixture to remove surface lipids and other contaminants, and air dried the samples. Feather vane material of about 300 μg encapsulated in tin capsules was then combusted using a Vario Isotope Cube Elementar Ananlyser (Elementar Analysensysteme GmbH, Germany). Resulting SO_2_, CO_2_ and N_2_ gases were transferred to the isotope ratio mass spectrometer (visION, Elementar Analysensysteme GmbH) via a continuous flow inlet system. Sample ^34^S/^32^S, ^13^C/^12^C and ^15^N/^14^N ratios are expressed in delta notation (*δ*^34^S, *δ*^13^C and *δ*^15^N) in parts per mil (‰) relative to V-PDB (Vienna-Pee Dee Belemnite) for δ^13^C, Air for δ^15^N, and V-CDT (Vienna-Canyon Diablo Troilite) for δ^34^S according to:

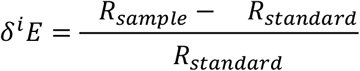

where the superscript *i* indicates the mass number of the heavier isotope of element *E* (*e.g*., ^34^S) and *R* the isotope ratio of *E* (*e.g*., ^34^S/^32^S) in the sample and in the internationally recognized standard. Values of δ^34^S, δ^13^C and δ^15^N were calculated against in-house standards which were calibrated against international reference materials using multi-point normalisation: fuel oil NBS-22 (IAEA International Atomic Energy Agency, Vienna, Austria; −30.03 ‰) and sugar IAEA-CH-6 (−10.45 ‰) for δ^13^C, L-glutamic acid USGS 40 (−26.39 ‰ and −4.5 ‰ for δ^13^C and δ^15^N), hair USGS 42 (δ^15^N = +8.05 ‰, δ^13^C = −21.09 ‰ and δ^34^S = +7.84 ± 0.25 ‰) and USGS 43 (δ^15^N = +8.44 ‰, δ^13^C = −21.28 ‰ and δ^34^S = +10.46 ± 0.22‰) for ^13^C/^12^C, ^15^N/^14^N and ^34^S/^32^S, barium sulphates IAEA-SO-5 and NBS 127 (IAEA) for ^34^S/^32^S. Each sample was analysed three times and we present its isotopic composition as an average value. Measurement error, calculated as one standard deviation of internal laboratory standard measured ten times, was ± 0.4‰ for *δ*^34^S, ± 0.1‰ for *δ*^13^C and ± 0.2‰ for *δ*^15^N. Stable isotope analyses were conducted in the Traceability Unit at the Research and Innovation Centre, Fondazione Edmund Mach, Italy.

### Covariates of δ^34^S

We gathered geographic distributions of potential atmospheric, anthropogenic, geographical and geological covariates of *δ*^34^S values, including SO_2_ atmospheric concentrations (Case and Krouse 1980, Ohizumi and Fukuzaki 1997), distance to coastline (Valenzuela et al. 2011, Zazzo et al. 2011), and bedrock age and type (Thode et al. 1953, Case and Krouse 1980). Moreover, we extracted the geographic distribution of NO_2_ atmospheric concentrations – tracing the intensity of various-source combustion – and log human population density, which affects the intensity of fossil fuel combustion (Garg et al. 2001, Martin et al. 2003). We also included surface elevation as a covariate, as we expected depletion in ^34^S with increasing elevation (atmospheric deposition), similar to the patterns observed in water isotopes (Dansgaard 1954, 1964). The original spatial resolution and sources of covariate distribution layers are detailed in Table 1. Geographic distribution layers were assembled using a cloud-based geospatial analysis platform Google Earth Engine (Gorelick et al. 2017).

**Table 1:** Covariates used for modelling the geographic distribution of feather δ^34^S values from Eurasian Reed Warblers across Europe.

| *Covariate* | *Units* | *Year* | *Original resolution* | *Source* |
| --- | --- | --- | --- | --- |
| SO<sub>2</sub> atmospheric concentration | mol m<sup>-2</sup> | 2018 | 1 km | (Copernicus Sentinel 2015a) |
| NO<sub>2</sub> atmospheric concentration | mol m<sup>-2</sup> | 2018 | 1 km | (Copernicus Sentinel 2015b) |
| Human density | log persons km<sup>-2</sup> | 2010 | 0.8 km | (CIESIN 2018) |
| Distance to coastline | km | – | 1 km | (NASA Ocean Biology Processing Group 2012) |
| Surface elevation | m a.s.l. | – | 0.8 km | (Danielson and Gesch 2011) |
| Bedrock type* | nominal (class) | – | discrete | (Pawlewicz et al. 2003) |
| Bedrock age* | nominal (years) | – | discrete | (Pawlewicz et al. 2003) |

*\*Layers were derived from vector layers with scale of 1:5000000.*

### Geostatistical analysis

We resampled the spatial resolution of all covariate layers at 2 km to match the scale of potential within-natal site movements of juvenile Eurasian Reed Warblers and extracted covariate values for each sample at each sampling location. We calculated average *δ*^34^S values at sampling sites or took the only measurement for further analysis (n = 69). To identify and quantify relationships between covariates and average feather *δ*^34^S at the sites, we employed a random forest regression model using the R package caret (Kuhn 2008). Random forest models are based on large number of regression trees grown using a randomly sampled subset of the training dataset (Breiman 2001, Belgiu and Dragut 2016). Random forest models can accommodate different data types including categorical covariates. This enabled us to predict abrupt changes in the spatial surfaces by avoiding smoothing over the regions (Bataille et al. 2018, 2021). We built the random forest model with 500 regression trees, randomly selecting two covariates at each node, and estimating model accuracy using leave-one-out cross-validation. In this process, each observation was withheld for subsequent validation of an entire random forest model, and this process was repeated for each observation (Vapnik 1998). We derived the relative importance of individual covariates (index of node purity) and visualised the partial dependencies (relationships) of these covariates on *δ*^34^S. We then applied the final random forest regression model to covariate values per grid cell and predicted the geographic distribution of feather *δ*^34^S values within the extent of prediction (Fig 2). We present the predicted *δ*^34^S isoscape for Europe <31°N, <61°N without Greece and Turkey, and for elevations up to 750 m a.s.l. These restrictions reflect the geographic range of sampling sites and their upper range of elevations (Fig 1).

**Fig 2.**
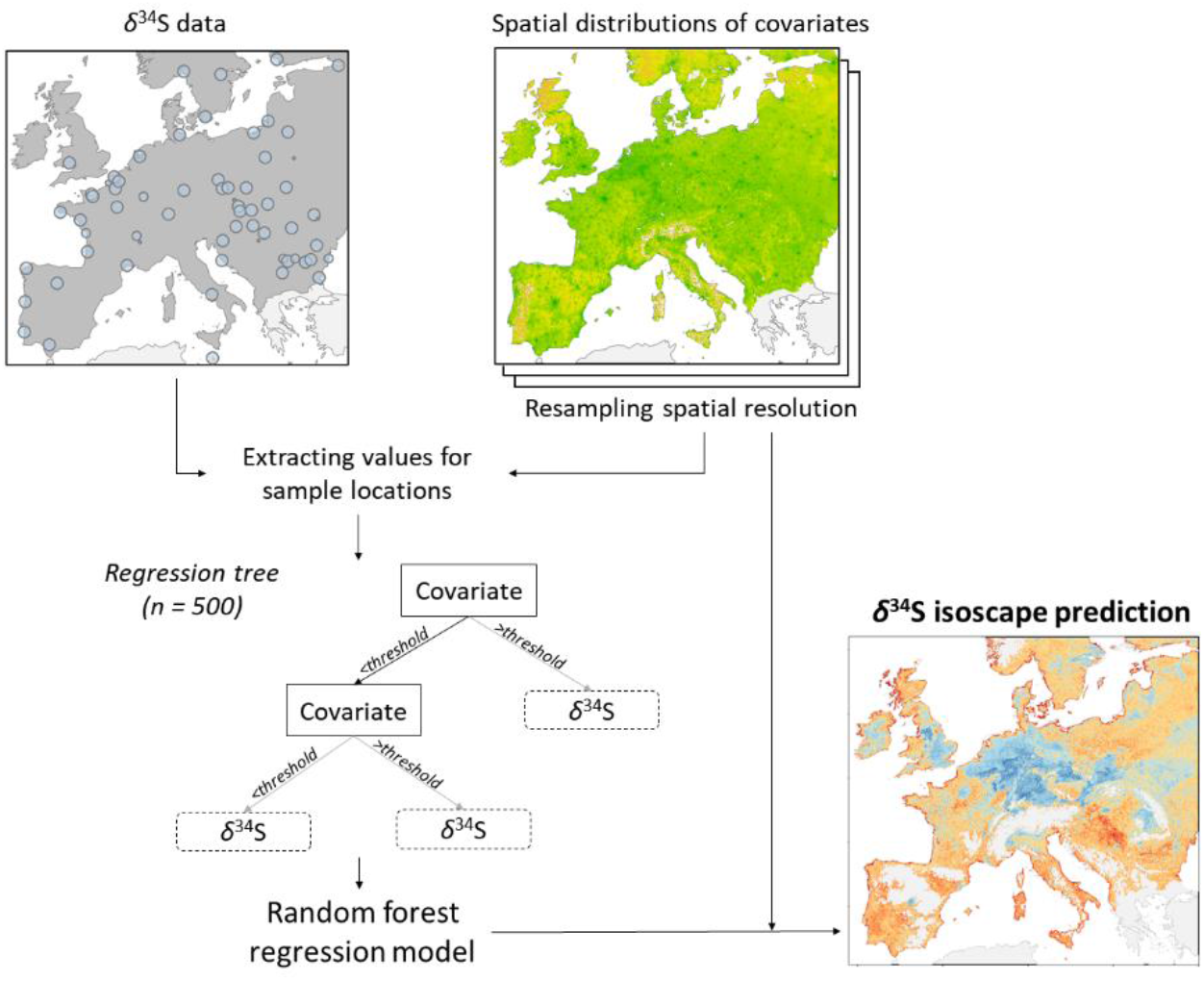
Flowchart showing the process of deriving the European feather δ^34^S isoscape. Dots in the map depict the geographic origin of samples used to derive the δ^34^S isoscape (n = 69). See Methods for more details.

Similarly, we quantified the relationships between the within-site standard deviation of *δ*^34^S values and the covariates. We calculated within-site standard deviation for each sampling site with at least three *δ*^34^S values (total of 230 samples from 59 sites) and employed the random forest regression model as detailed above. Finally, we quantified the relationship between average within-site *δ*^34^S values and within-site standard deviations of *δ*^34^S values using Pearson’s correlation coefficients.

### Complementarity of δ^34^S with other isotopes

We assessed the complementarity of *δ*^34^S on individual and isoscape levels. On the individual level, we quantified Pearson’s correlations between feather *δ*^34^S values, and the feather *δ*^13^C and *δ*^15^N values measured in Eurasian Reed Warblers (n = 242). Similarly, we assessed isoscape-level complementarity to identify similarities among spatial patterns of isoscapes. We compared spatial patterns using per-grid-cell Pearson’s correlation coefficients between the predicted *δ*^34^S isoscape and the previously known feather-based *δ*^2^H and *δ*^13^C isoscapes. At both individual and isoscape levels low correlations indicate high level of complementarity.

For assessing the complementarity on the isoscape level, we prepared the European feather *δ*^2^H isoscape. We extracted *δ*^2^H in precipitation from Global Network of Isotopes in Precipitation and applied a geospatial model to predict the *δ*^2^H isoscape using precipitation, average temperature, surface elevation, absolute latitude as independent explanatory variables via a web-based computing platform IsoMAP (Bowen et al. 2014; task key: 84119). We calibrated the resulting isoscape of *δ*^2^H in precipitation using the equation *δ*^2^H_feather_ = 1.28 × *δ*^2^H_precipitation_ −10.29 to account for discrimination between precipitation and feather *δ*^2^H in the Eurasian Reed Warbler derived by Procházka et al. (2013), and resampled to the spatial resolution the predicted feather *δ*^34^S isoscape.

Similarly, we prepared the European plant *δ*^13^C isoscape by combining climatic (Abatzoglou et al. 2018), land cover (Buchhorn et al. 2020) and crop production layers (Yu et al. 2020). We adapted the process described in Powell et al. (2012) and Powell et al. (2019) using Google Earth Engine (Gorelick et al. 2017). We accounted for an assumed plant–feather discrimination by adding +2‰ to the plant *δ*^13^C isoscape (Hobson et al. 2012) and resampled to the spatial resolution of the predicted feather *δ*^34^S isoscape. Analyses were conducted in R 4.1.2 (R Core Team 2021).

## Results

### Geographic distribution of δ^34^S and main covariates

The predicted feather *δ*^34^S isoscape features a region depleted in ^34^S over eastern Great Britain, Benelux, Germany, Poland, and northern Ukraine. In contrast, northern Europe (i.e., Norway, Denmark, Sweden, Finland, and the Baltic states) and western to southern Europe (i.e., the Iberian Peninsula, France, Italy, the Balkans, Romania, and Bulgaria), together with the European coastline were more enriched in ^34^S (Fig 3).

**Fig 3.**
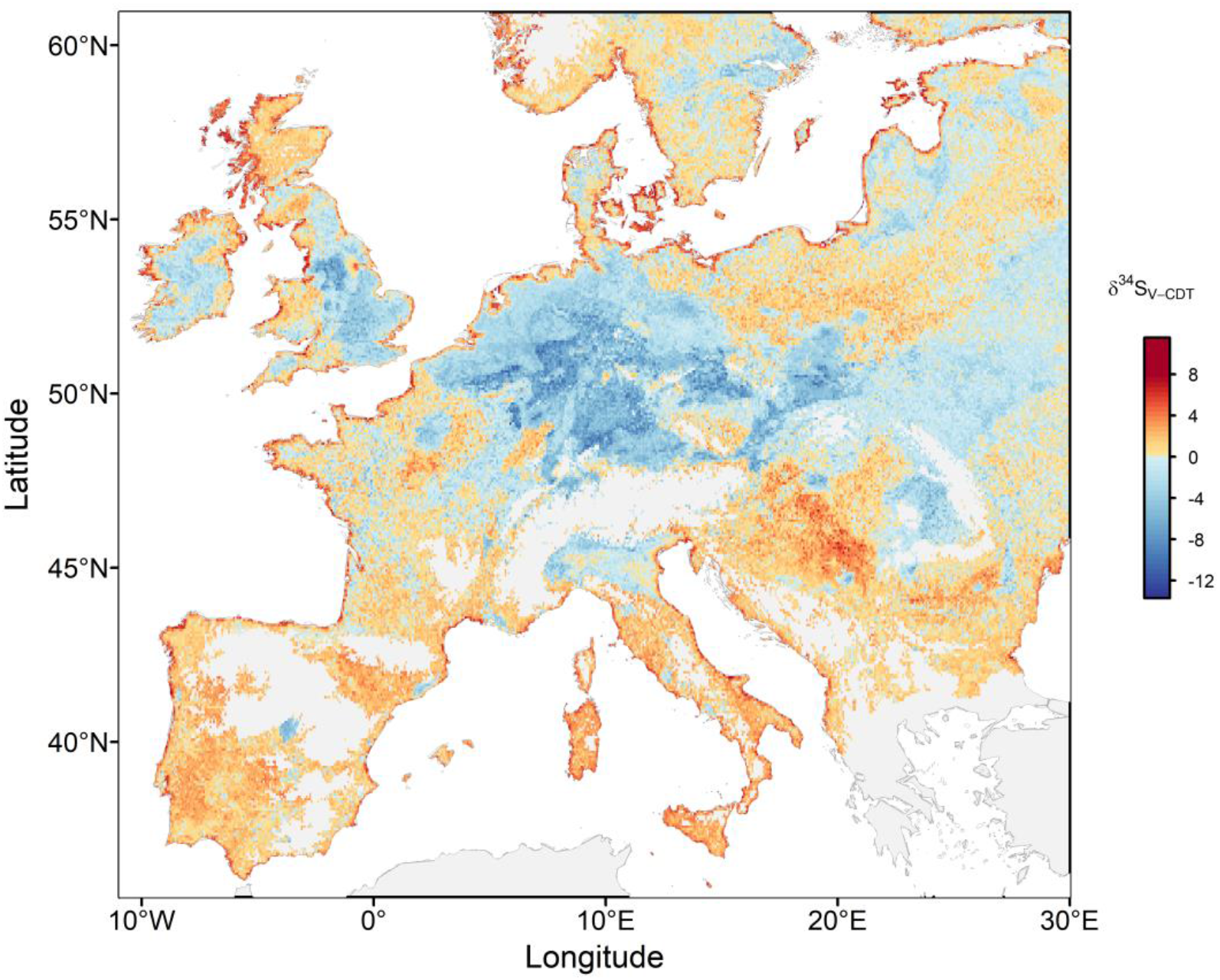
The predicted geographic distribution of δ^34^S values in Europe derived using Eurasian Reed Warbler feather samples of known spatial origin, grey colour represents regions outside prediction range.

The most important covariates of *δ*^34^S values were distance to coastline, surface elevation, and atmospheric concentrations of SO_2_ (Fig 4). The relationships between the covariates and the *δ*^34^S values were (i) negative (depletion in ^34^S) for distance to coastline, surface elevation and human density, (ii) positive (enrichment in ^34^S) for bedrock age, and (iii) more complex for atmospheric concentrations of SO_2_ and NO^2^ (Fig 4).

**Fig 4.**
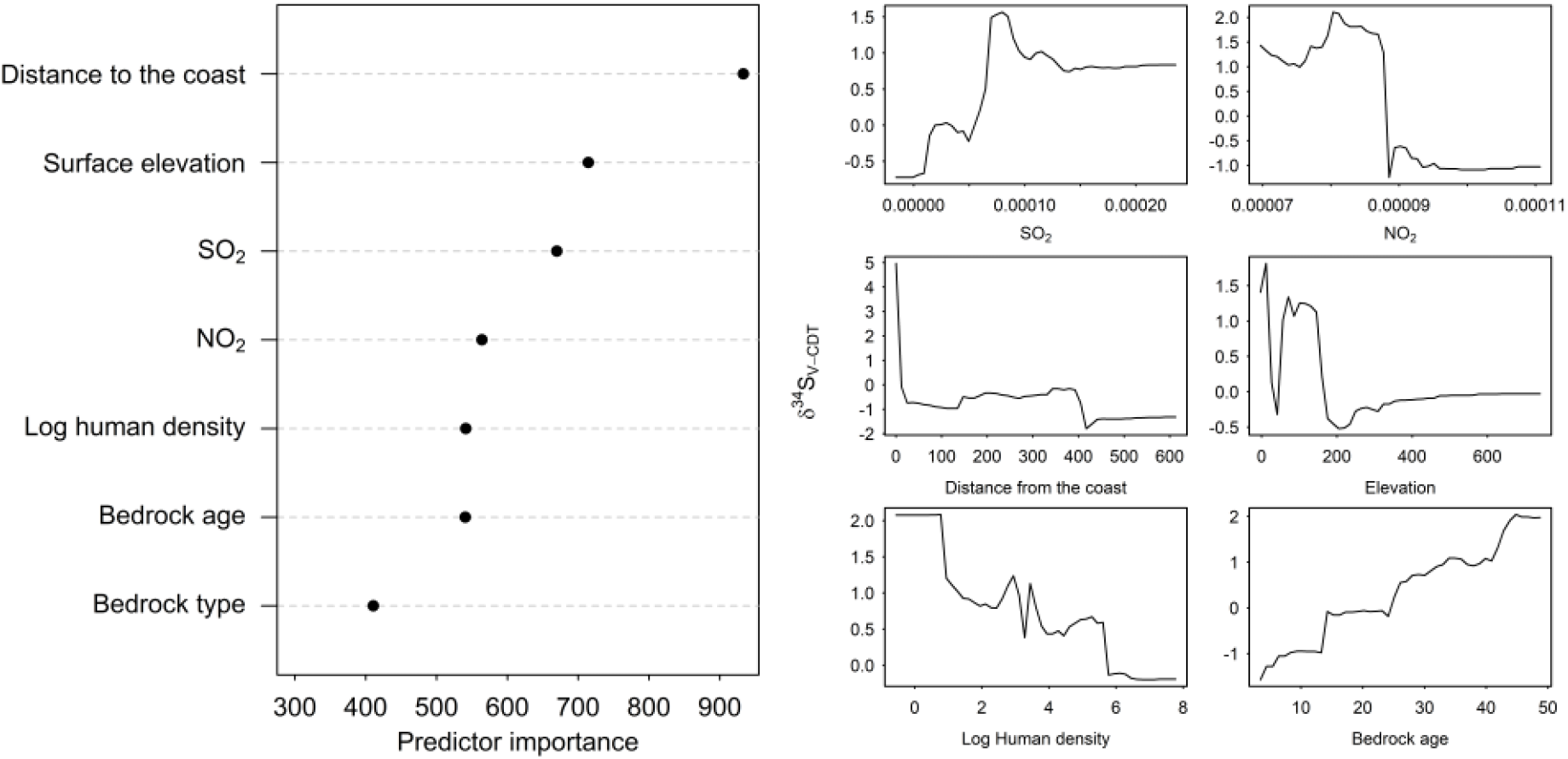
Relative importance of covariates (left) and partial dependencies of δ^34^S on covariates (right).

Overall, the random forest regression model explained 90% of the variance in the training dataset (predicted vs. observed values; slope = 1.5 [SE = 0.1], intercept = −0.4 [0.3]) and the cross-validation explained 15% of variance (slope = 0.8 [0.3], intercept = 0.4 [1.0]). Root mean square error of the final model was 6.1‰ representing 14% of the range of the *δ*^34^S in the training dataset.

### Within-site variation in δ^34^S

The average within-site variation (SD) in feather *δ*^34^S was 2.1 ‰ (median = 1.3, min = 0.1, 1^st^ quartile = 0.7, 3^rd^ quartile = 2.5, max = 9.1). Overall, the highest within-site variation was detected in some inland parts of Europe (i.e., France, Germany, Hungary, southern Poland) while the lowest values appeared in coastal regions (Fig 5). The correlation between the within-site averages and within-site standard deviations was −0.21 (Pearson’s correlation; 95% CI [-0.44, 0.05]) and sites close to the coastline tended to show low within-site variability (ρ = 0.29 95% CI [0.05, 0.51]). The random forest regression model of within-site variability explained 87% of variance in the training dataset (slope = 2.3 [SE = 0.1], intercept = −2.2 [0.2]) but the cross-validation explained no variance (slope = 0.3 [SE = 0.4], intercept = 1.4 [SE = 0.9]). Root mean square error of the model was 1.6‰ representing 18% of the range of the values in the training dataset. Due to the low performance of the model, we did not predict the geographic distribution of within-site variability across Europe and only present raw values (Fig 5).

**Fig 5.**
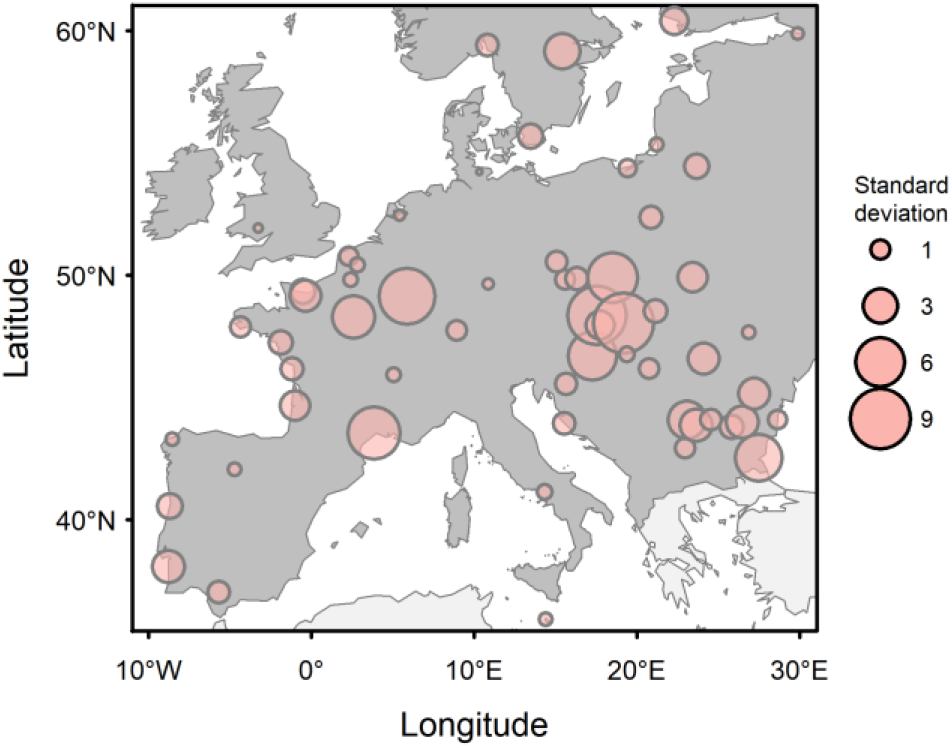
Within-site variability presented as standard deviations of feather δ^34^S values at sites with at least three samples collected (59 sites).

### Complementarity of δ^34^S with other isotopes

The feather *δ*^34^S values ranged between −27.6 and 20.4‰ (mean = 1.0, SD = 8.5, n = 242), *δ*^13^C values ranged between −28.9 and −11.5‰ (mean = −23.9, SD = 2.1, n = 242) and *δ*^15^N values ranged between 6.5 and 25.6‰ (mean = 12.6, SD = 2.7, n = 242). The feather *δ*^34^S values were weakly related to both *δ*^13^C (ρ = 0.29; 95% CI [0.17, 0.40]) and *δ*^15^N (ρ = −0.19; 95% CI [-0.30, −0.06]; Fig. 6). Similarly, we identified only weak relationships of the predicted feather European *δ*^34^S isoscape with the feather *δ*^2^H (ρ = 0.19) or feather *δ*^13^C isoscapes (ρ = 0.01; Fig. 7).

**Fig 6.**
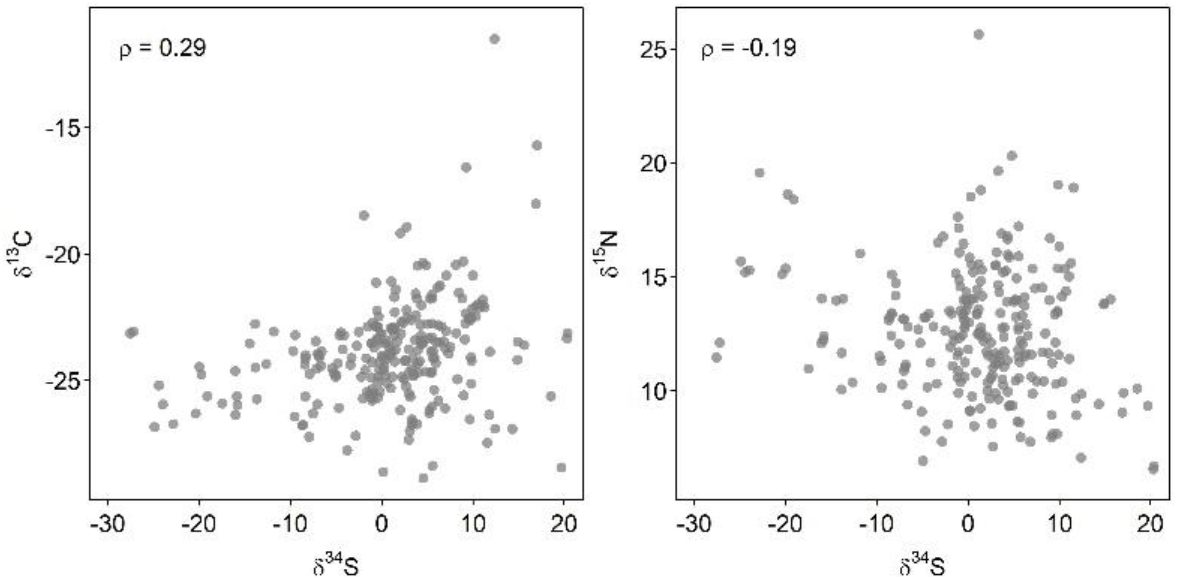
Relationship between feather δ^34^S and δ^13^C (left) and δ^34^S and δ^15^N (right) measurements (n = 242) including Pearson’s correlation coefficients.

**Fig 7.**
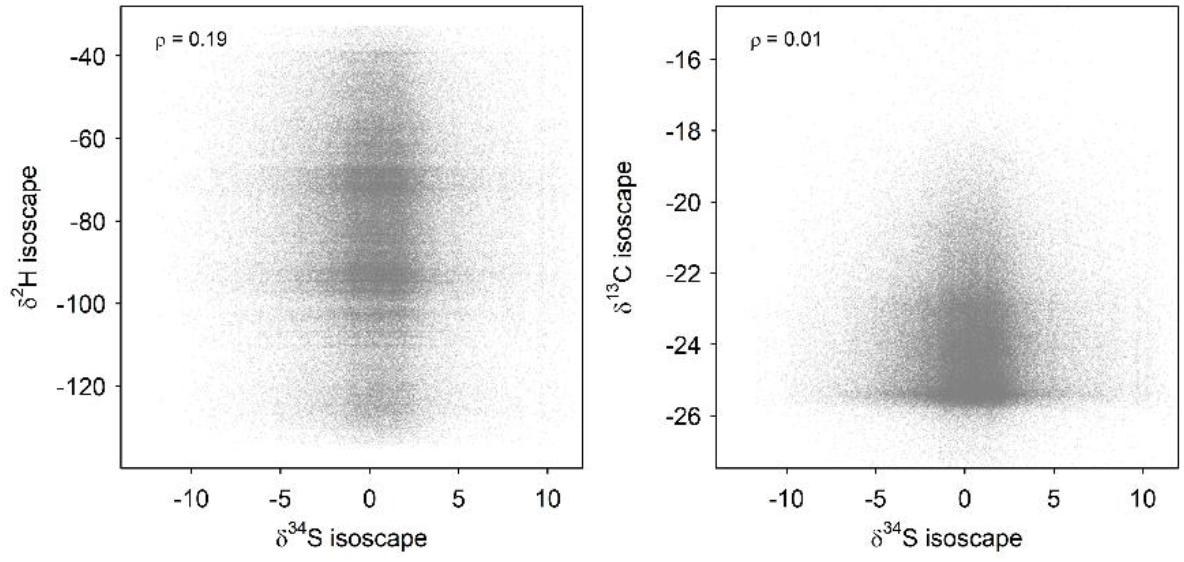
Relationships between the feather δ^34^S and δ^2^H isoscapes (left), and the feather δ^34^S and δ^13^C isoscapes (right) including Pearson’s correlation coefficients.

## Discussion

We predicted the geographic distribution of *δ*^34^S values in Europe using feather samples of known geographical origin. The *δ*^34^S isoscape shows high *δ*^34^S values along the European coast and only slight differences in *δ*^34^S values between the central and southern parts of Europe. The *δ*^34^S values also varied highly within sites, especially, in inland parts of Europe. In the absence of a systematic spatial pattern, the *δ*^34^S isoscape, derived from feathers grown in wetlands, appears to be of limited use alone. However, its high level of complemetarity with other isoscapes suggests combining multiple isoscapes to increase animal tracing precision.

### Geographic distribution and main covariates of δ^34^S

Coastal regions being the most enriched in ^34^S and the distance to coastline being a key covariate of the geographic distribution of *δ*^34^S is in line with our predictions and previous studies (Fry 2006, Zazzo et al. 2011, Amrani et al. 2013). In contrast, the inland parts of Europe showed only slight differences in *δ*^34^S values between the regions and the absence of a systematic spatial pattern thus contrasts with previous regional- and continental-scales assessments (Wadleigh and Blake 1999, Valenzuela et al. 2011, Bataille et al. 2021, Newton 2021; Brlík et al. 2022a). Among the explanations for the absence of such systematic spatial patterns in Europe could be: (i) the complex geography of the European continent, (ii) the spatial clustering of human population with related distribution of fossil fuel combustion intensity, and (iii) the effect of variability in redox state of wetland soils that influences tissue *δ*^34^S isotopic compositions.

The transportation of marine sulphate aerosols creates the coast–inland gradient of *δ*^34^S values on regional and continental scales (Valenzuela et al. 2011, Zazzo et al. 2011). However, the European continent has complex coastal geography, intersected by mountain ranges, and contains regions with distinct climates (Beck et al. 2018) and variable patterns (Troen and Petersen 1989). Complex geography and wind systems could thus disrupt inland transportation of marine sulphates, partly explaining the patchiness of the predicted *δ*^34^S isoscape. This explanation would be indirectly supported by the presence of a *δ*^34^S gradient over homogeneous land masses in North America (Valenzuela et al. 2011) or sub-Saharan Africa (Procházka et al. 2018; Brlík et al. 2022a). However, our explanation lacks direct support and contradicts the recent European *δ*^34^S isoscape althought based on mostly archaeological and historical collagen samples that shows a continent-wide coast–inland gradient of *δ*^34^S values within Europe (Bataille et al. 2021).

Human populations have boomed in the past century as well as the intensity of anthropogenic activities. Human population growth has influenced atmospheric concentrations of combustion gases (i.e., SO_2_ and NO_2_) that we identified as important covariates of *δ*^34^S values. Human population and related anthropogenic sources of these gases are, however, spatially clustered with numerous local peaks (e.g., in urban agglomerations and around factories) with highest densities in central and western parts of Europe. A clustered geographic distribution of these covariates could further disrupt coast–inland gradient of *δ*^34^S values caused by marine-derived sulphate aerosols (Rees et al. 1978, Zazzo et al. 2011) and result in patchiness of the predicted feather *δ*^34^S isoscape.

The overall patchiness of the predicted feather *δ*^34^S isoscape could be related to biological conditions in wetlands – the breeding habitat of the Eurasian Reed Warbler and the only habitat we sampled. Wetlands store water, and accumulate organic material and nutrients from surrounding areas. Consequently, geochemical conditions in the wetland may be affected by the catchment capacity of water reservoirs, collecting water leaching various bedrocks in the surrounding region. Moreover, high availability of nutrients and low oxygen levels support development of anaerobic conditions (Cherry 2011) and associated bacterial communities. These bacteria affect *δ*^34^S composition through fractionation against ^34^S, resulting in substantial depletion in ^34^S in solid-phase sulfur-bearing minerals such as pyrite (Chambers and Trudinger 1979, Thode 1991). Unfortunately, the anaerobic conditions in wetlands cannot be predicted from remote sensing imagery and are thus difficult to quantify on continental scales. Finally, the predicted feather *δ*^34^S isoscape considers the relationships identified within wetland sampling sites but applies these relationships to predict *δ*^34^S values over Europe potentially leading to bias (due to e.g., an absence of low redox conditions in terrestrial habitats).

### Within-site variation in δ^34^S

We found large differences in within-site *δ*^34^S variation with the highest values detected at some inland sites in Europe. In contrast, the coastal sites showed mostly low variation, which could be explained by the prevailing impact of marine-derived sulphates with uniform *δ*^34^S values (Rees et al. 1978). Within-site variation in *δ*^34^S also tended to increase with increasing distance to coastline likely due to reduced transport of marine sulphates and an increasing impact of low redox conditions in wetland environments (Chambers and Trudinger 1979, Thode 1991), which could vary locally both between patches within an individual wetland and regionally among wetlands.

Moreover, our measurements of within-site variability are based on three to four feather samples collected from juvenile Eurasian Reed Warblers during the post-fledgling period at each site. The limited sample sizes could have contributed to large within-site variation due to stochasticity. Potential post-fledgling dispersals of juvenile Eurasian Reed Warblers between wetlands (Mukhin 2004) could further increase the variation detected at sampling sites. Similarly, the fledglings or adults feeding them before fledgling could opportunistically use food resources within the wetland as well as in the surrounding terrestrial habitats with different isotopic signatures adding to the within-site variation in *δ*^34^S (Catchpole 1971, Król 1984).

### Complementarity of δ^34^S with other isotopes

Our results showed high complementarity of *δ*^34^S with other isotopes on both the individual (feather samples) and spatial (isoscape) levels suggesting different environmental drivers differ among isotopes. However, the high isoscape-level complementarity should be interpreted with caution. The feather samples used to predict the *δ*^34^S isoscape in this study were formed in a single habitat type – wetlands. Wetland conditions likely differ from terrestrial conditions as indicated by the differences in geographic pattern of our isoscape and the European *δ*^34^S isoscape from archaeological remains (Bataille et al. 2021). Nevertheless, our results show a high potential of *δ*^34^S values in Europe, especially, in combination with other isotopes but a dense sampling of tissues from diverse taxa and across habitats is needed.

## Conclusions

The predicted geographic distribution of *δ*^34^S values in Europe shows high *δ*^34^S values along the coastline in Europe and slight differences in *δ*^34^S values between the central and southern parts of Europe. The absence of a systematic spatial pattern and high within-site variation limits the sole use of the *δ*^34^S isoscape. However, high levels of complementarity on the individual and isoscape levels suggest the potential of combining isoscapes to increase precision of animal tracing. Therefore, we encourage future studies to focus on geographic distribution of *δ*^34^S in homogeneous and large landmasses in Africa, South America or East Asia. Moreover, we recommend to predict geographic distributions of *δ*^34^S variables from samples collected in various environments, habitats, and from different animal species. We believe these efforts will significantly advance the knowledge on the *δ*^34^S geographic distributions and enhance their applications in ecological and forensic studies.

## Acknowledgments

We are obliged to following bird ringers who helped us collect feathers samples in Europe: A. Leprêtre, A. Ożarowska, A. Trnka, A. Sponga, A. Pranaitis, B. Vollot, B. Fontaine, B. Bargain, C. Coleiro, R. Galea, C. Z. Martínez, C. Heroguel, C. Ion, D. Vigour, D. Dimitrov, M. Ilieva, D. Ragyov, D. Zhuravlev, D. Leoke, E. Baltag, F. G. Lopez, G. Goujon, J. R. Álvarez, J. Hlaváček, J. J. Baptiste, J. Kralj, J. Reif, J. Beier, J. M. Neto, K. and F. Breek, K. Bedev, K. Bairaktaridau, L. F. P. da Silva, M. Haluzík, M. Šćiban, M. Leconte, M. Olekšák, O. Zakala, P. S. Ranke, P. Matyjasiak, R. Thomas, R. Patapavičius, R. Slobodník, S. Gautier, S. Scebba, S. Capasso, S. Bräger, V. Cohez, V. Fedorov, V. Jusys, W. Fiedler, X.

Commecy, Y. Beauvallet and Z. Karcza. We thank Gabriel Bowen and the Isotopes in Spatial Ecology and Biogeochemistry short course held at the University of Utah for providing the stimulating environment that led to this project. We are also thankful to Carol Kendall for initial advice and reviewers for their comments.

## Authors Contributions

VB conceived the idea of the study. VB, FJ, GO and PP collected or provided samples. LB and FC analysed stable isotopic composition of samples. VB designed the methodology with input from RLP, PP, CAS and MBW. VB conducted the analyses with input from MBW and RLP and took lead in the manuscript writing. All authors read, commented, and approved the final version of the manuscript.

## Conflict of Interest Statement

Authors declare no conflict of interest.

## Funding

This work was supported by the Grant Agency of Charles University (grant no. 254119), the National Science Foundation (grant no. DBI-1565128) and the Czech Science Foundation (grant no. 20-00648S).

## Notes

https://doi.org/10.5281/zenodo.7315567

### Competing Interest Statement

The authors have declared no competing interest.

